# An IBD-associated pathobiont synergises with NSAID to promote colitis which is blocked by NLRP3 inflammasome and Caspase-8 inhibitors

**DOI:** 10.1101/2022.08.18.504384

**Authors:** Raminder Singh, Valerio Rossini, Stephen R. Stockdale, Gonzalo Saiz-Gonzalo, Naomi Hanrahan, Tanya D’ Souza, Adam Clooney, Lorraine A. Draper, Colin Hill, Ken Nally, Fergus Shanahan, Stefan Andersson-Engels, Silvia Melgar

## Abstract

Conflicting evidence exists on the association between consumption of non-steroidal anti-inflammatory drugs (NSAIDs) and symptomatic worsening of inflammatory bowel disease (IBD). We hypothesise that the heterogeneous prevalence of pathobionts [e.g., adherent-invasive *Escherichia* coli (AIEC)], might explain this inconsistent NSAIDs/IBD correlation. Using *IL10* ^*-/-*^ mice, we show aggravation of colitis in AIEC-colonised animals fed NSAID. This is accompanied by activation of the NLRP3 inflammasome, Caspase-8, apoptosis and pyroptosis; features not seen in mice exposed to AIEC or NSAID alone, revealing an AIEC/NSAID synergistic effect. Inhibition of NLRP3 or Caspase-8 activity ameliorated colitis, with reduction in NLRP3 inflammasome activation, cell death markers and activated T-cells and macrophages, improved histology and increased abundance of *Clostridium* cluster XIVa species. Our findings provide mechanistic insights into how NSAID and an opportunistic gut-pathobiont can synergise to worsen IBD symptoms. Thus, targeting the NLRP3 inflammasome and Caspase-8 could be a potential therapeutic strategy in patients with NSAID-worsened inflammation.

## INTRODUCTION

Inflammatory bowel disease (IBD) is a chronic inflammatory state of the gastrointestinal tract, including Crohn’s disease (CD) and Ulcerative colitis (UC). The aetiology of IBD is unknown, but the collective evidence indicates that a dysregulated immune response to commensal enteric microorganisms (one of the hallmarks of IBD), leads to uncontrolled chronic inflammation in the genetically susceptible host (Round and Mazmanian, 2009)

Non-steroidal anti-inflammatory drugs (NSAIDs) are commonly used for various inflammatory conditions, including IBD-associated extraintestinal manifestations. But mucosal injury and ulcers are common adverse effects of NSAID use and are therefore believed to exacerbate inflammation in patients with IBD (Ananthakrishnan et al., 2012). However, the current literature on the association between NSAIDs and IBD is inconclusive (Klein and Eliakim, 2010, Moninuola et al., 2018). Evidence to date suggests that the gut microbiota might play a critical role in NSAID-promoted inflammation (Wang et al., 2021). For example, germ-free rats are resistant to NSAID-induced intestinal damage (Robert and Asano, 1977) and an increase in the relative-abundance of potentially harming bacteria such as *Rikenellaceae, Pseudomonadaceae, Propionibacteriaceae*, and *Puniceicoccaceae* was observed in individuals taking ibuprofen (Rogers and Aronoff, 2016). In addition, consumption of a *Bifidobacterium* strain reduced NSAID-induced ulceration in healthy volunteers (Mortensen et al., 2019).

It is possible that genetic background and environmental factors can contribute to disease triggered by pathogenic commensals, so-called pathobionts (Chow et al., 2011). Adherent-invasive *Escherichia* coli (AIEC) is a gut pathobiont widely prevalent in IBD patients in Western countries, up to 10% and 62% in patients with UC and CD, respectively (Nadalian et al., 2021, Palmela et al., 2018). AIEC has been reported to trigger inflammation in genetically predisposed mice and can promote fibrosis in chemically induced colitis (Small et al., 2013). A recent study reported that AIEC might be associated with the early phase of recurrence in patients with CD (Buisson et al., 2022), further indicating their possible contribution to CD pathogenesis.

NOD-like receptors (NLRs) are an important family of pathogen recognition receptors (PRRs) that sense pathogen and commensal associated molecular patterns (PAMPs), or danger associated molecular patterns (DAMPs) leading to co-ordinated innate and adaptive immune responses. One of the most intensely studied members of this NLR family is NLRP3, whose activation leads to the formation of the NLRP3 multiprotein inflammasome complex. Formation of this complex activates caspase-1 and subsequent cleavage of downstream substrates such as the effector inflammatory cytokines, pro-IL1β and pro-IL18 (Broz and Dixit, 2016). Activation of NLRP3 has been reported in pre-clinical models of intestinal infection and colitis, in human IBD tissue and by AIEC in macrophages (Aguilera et al., 2014, De la Fuente et al., 2014). Genome-wide association studies have identified a SNP in the region of the NLRP3 gene possibly contributing to disease susceptibility in CD, and a recent study reported a loss-of-function mutation in the CARD8 domain, causing NLRP3 activation and CD (Mao et al., 2018). NLRP3 has also been shown to play a crucial role in NSAID-induced enteropathy (Higashimori et al., 2016).

One of the hallmarks of chronic inflammation is the increased death (apoptosis) of intestinal epithelial cells in patients with IBD (Di Sabatino et al., 2003). One initiator caspase which acts as a central regulator of the crosstalk and plasticity between multiple cell death pathways (apoptosis, necroptosis and pyroptosis) and inflammation, is Caspase-8 (Newton et al., 2021). Various studies have shown the role of Caspase-8 in intestinal inflammation (Lehle et al., 2019, Gunther et al., 2011). Activation of death receptors (e.g., TNFR), Toll-like receptors (TLR3/-4), the z-form nucleic acid sensor (ZBP1), or the intracellular RNA sensor RIG-I leads to the recruitment of Caspase-8 to the intracellular signalling complex, followed by its autoproteolytic cleavage and activation. Active caspase-8 cleaves its substrate caspase-3/7, initiating apoptosis. In addition, active caspase-3/7 can also cleave poly (ADP-ribose) polymerase (PARP), resulting in aberrant DNA repair and apoptosis (Mashimo et al., 2021, Newton et al., 2021). Active Caspase-8 inhibits necroptosis by supressing the function of receptor-interacting protein kinase 1 (RIPK1) and RIPK3, to phosphorylate the pore-forming mixed lineage kinase domain-like (MLKL) protein (Bergsbaken et al., 2009).

Another cell death mechanism provoked by inflammation and microbial infections, e.g., *Salmonella*, is pyroptosis. This cell death can be induced by the canonical activation of Caspase-1 or via the non-canonical caspases, Caspase-11 (mice) and Caspase-4/5 (humans) (Bergsbaken et al., 2009), leading to the cleavage of its main executor gasdermin D (GSDMD). Interestingly, Caspase-8 has been shown to cleave GSDMD leading to pyroptosis (Gram et al., 2019). Caspase-8 can act both upstream and downstream of NLRP3 inflammasome activation. On one hand, it triggers NLRP3 activation via both its enzymatic activity and scaffolding function (Sarhan et al., 2018, Kang et al., 2015, Chi et al., 2014, Antonopoulos et al., 2015). On the other hand, NLRP3 can regulate Caspase-8 activation in epithelial cells independent of Caspase-1 activation or cytokine production (Chung et al., 2016).

Based on the collected evidence to date, we hypothesised that the apparent inconsistency in reports regarding the impact of NSAIDs on IBD disease activity could be explained by the absence/presence of AIEC in these patient cohorts. In this study, we used interleukin-10 deficient (*IL10* ^*-/-*^) mice - a pre-clinical model of IBD - to test our hypothesis. We show that AIEC collaborates with NSAID to induce colitis and cell death via the NLRP3 inflammasome with the contribution from Caspase-8 to this overall phenotype. This provides mechanistic insight into how NSAIDs in the presence of an opportunistic gut pathobiont could worsen IBD symptoms.

## RESULTS

### AIEC-mediated Sensitization of IL10 ^-/-^ mice to Piroxicam induced-inflammation is associated with activation of the inflammasome, apoptosis and pyroptosis

To test our hypothesis, we examined the ability of AIEC-HM605, a strain of AIEC isolated from colonic biopsies of a patient with CD (Martin et al., 2004), to sensitise *IL10* ^*-/-*^ mice to NSAID (piroxicam) induced inflammation. We developed a model whereby initial pre-treatment to *IL10* ^*-/-*^ mice with streptomycin facilitated colonisation with AIEC (Barthel et al., 2003) (Figure S1A). Subsequent feeding of these mice with 100 ppm of piroxicam in regular chow for 5 days followed by 9 days of normal chow resulted in colitis (Figure 1A). The initial colonisation of AIEC dropped from 10 ^9^ to 10 ^4^ cfu/g at day 7, followed by a stable AIEC colonisation throughout the study (Figure S1A). In line with other models of colitis (Melgar et al., 2005a), only the mice colonised with AIEC and subsequently treated with NSAID had shorter and heavier colons compared to animals exposed to piroxicam or AIEC alone (Figure 1B). This was accompanied by a loss of colonic epithelial integrity indicated by the significant reduction in expression of colonic *Zo1* and *Muc2* genes (Figure 1C) and a significant induction in the expression of inflammatory genes and proteins (*Il17a, Cxcl2, Ifng* and IL1β, IFNγ and mKC, Figure 1D-E, S1B-C).

**Figure 1.**
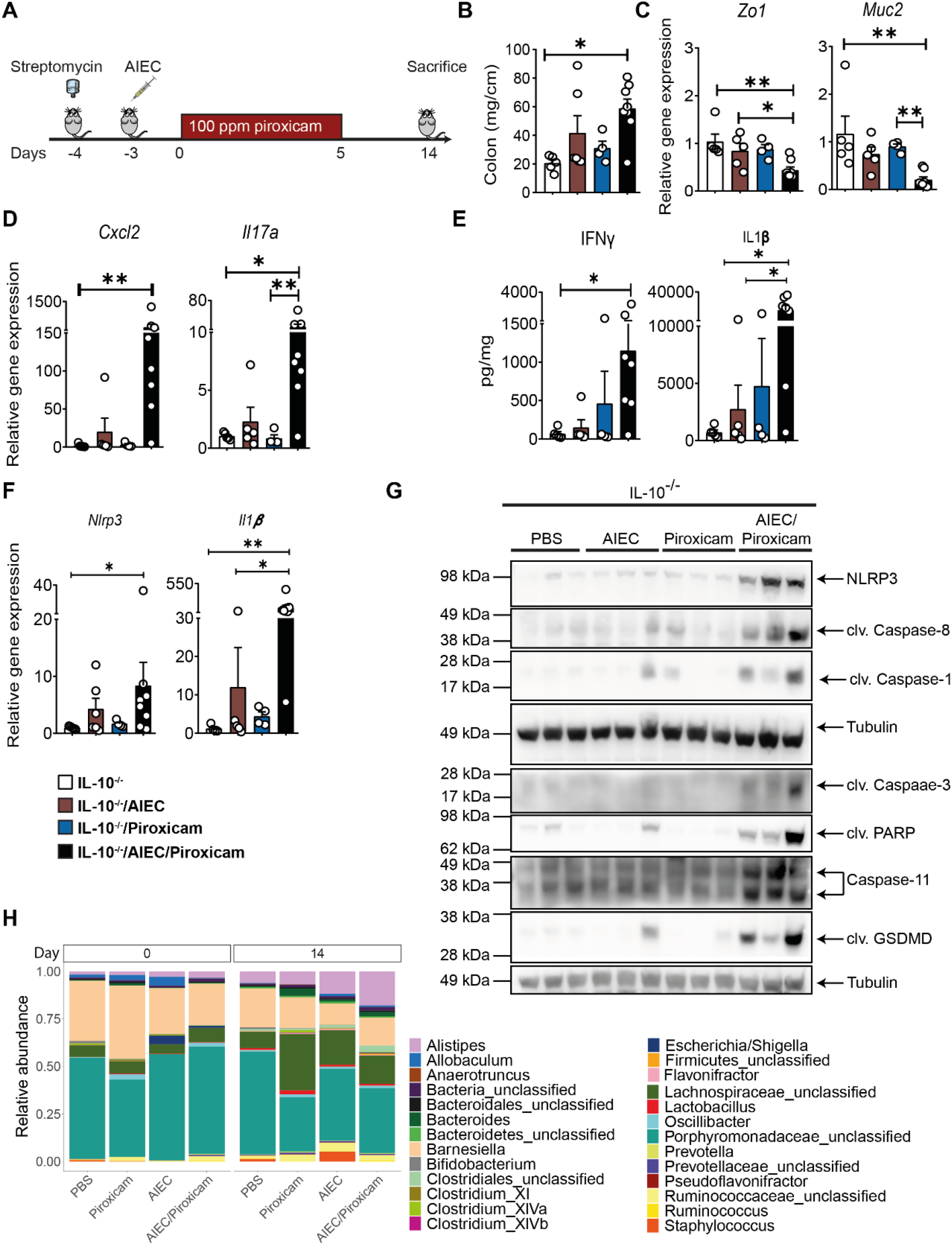
IL-10 ^-/-^ mice colonised with AIEC and fed Piroxicam develop colitis associated with activation of Nlrp3 inflammasome and Caspase-8. (A) Study design. IL-10 ^-/-^ mice were given streptomycin (5 g/L) in drinking water *ad libitum* for 24 hrs, followed by oral gavaged with approx. 10 ^9^ AIEC colony forming units (CFU), 100ppm of piroxicam homogenised in normal chow for 5 days followed by normal chow until day 14. (B) Distal colon weight. (C) Real-time (RT)-qPCR for colonic epithelial and (D) inflammatory markers. (E) Protein levels of colonic inflammatory cytokines and chemokines. (F) RT-qPCR for colonic *Nlrp3* and *Il1b*. (G) Western blot of markers of NLRP3 inflammasome, Caspase-8, apoptosis and pyroptosis markers. (H) Faecal bacteria relative-abundance at genus-level. Microbiota analysis, n=4-9/group and n=3-8/group for RT-qPCR, protein, western blot and necroscopy. Data are presented as mean ± SEM. Significance was determined using one-way ANOVA with Bonferroni or Kruskal-Wallis test with Dunn’s multiple comparison test, respectively. * p < 0.05; ** p <0.01; *** p <0.001.

Because the NLRP3 inflammasome has previously been shown to be activated by AIEC strains *in vitro* and in NSAID enteropathy (De la Fuente et al., 2014, Higashimori et al., 2016), we analysed its activation by AIEC/NSAID in our model. Indeed, a significant upregulation of colonic NLRP3, caspase-1 and its downstream target IL-1β was revealed at the gene expression and protein level in the AIEC/piroxicam combination group but not in mice colonised only with AIEC or piroxicam alone (Figure 1E-G). We also observed activation of Caspase-8 in the AIEC/piroxicam combination group (Figure 1G). To confirm Caspase-8 activation, we assayed for the protein expression of its downstream target NEDD4-binding protein 1 (N4BP1) (Gitlin et al., 2020). Indeed, cleaved N4BP1 was only detected in AIEC/piroxicam combination group (Figure S1D). Since Caspase-8 can control apoptosis by activating Caspase-3 by cleaving it (Kim et al., 2020), we assayed for its cleaved expression and found it to be highly expressed in AIEC/piroxicam group when compared to AIEC or piroxicam only groups (Figure 1G). A similar cleavage in PARP was also observed in the AIEC/piroxicam group (Figure 1G). No activation of necroptosis, examined by protein levels of MLKL or RIPK1, was seen in any of the groups (data not shown). When we examined the expression of the main executor marker of pyroptosis, cleaved GSDMD, we found it only expressed in the AIEC/piroxicam group (Figure 1G), indicating a synergistic activation of AIEC/piroxicam of pyroptosis.

Collectively, these observations validate our initial hypothesis indicating that AIEC colonisation in a genetically predisposed host renders them susceptible to NSAID-induced inflammation. This inflammation is accompanied by activation of the inflammasome, apoptosis and pyroptosis.

### Colonisation with AIEC and Piroxicam treatment provoke modest alterations of the gut microbiome in IL10 ^-/-^ mice

To determine the impact of piroxicam and/or AIEC exposure on the gut microbiome, we conducted 16S rRNA gene analysis on faecal samples collected at d0 and 14 (Figure 1A). Across treatment groups, an increase in intra-sample alpha-diversity was observed over time (Figure S1F), suggesting a gradual recovery of mice microbiomes following Streptomycin treatment. While there is a significant (p < 0.05) increase in alpha-diversity observed in AIEC, and AIEC/Piroxicam treated mice between day 0 and day 14 (Figure S1F), this trend was non-significant after false-discovery rate (fdr) correction.

Visually, the ordination of mice 16S rRNA composition shows beta-diversity differences are greater in mice colonised only with AIEC and/or treated with Piroxicam compared to PBS-treated controls (Figure S1G). Permutational multivariate analysis of variance (PERMANOVA) tests were performed on inter-sample beta-diversity distances to discern the variables significantly contributing to data dispersion (i.e., variance). This analysis revealed that the days post piroxicam variable accounted for 14.9% of the 16S rRNA data variance (Figure S1G).

The compositional bar plots on 16S rRNA data show only taxa with a total aggregated relative abundance across all mice greater than 0.1% (Figure 1H). As expected, *Escherichia/Shigella* are detected at day 0 (i.e., 4 days post-AIEC colonisation) of the relevant treated animals. However, significant differences in other individual 16S microbial taxa were not observed between mice over time after fdr correction, although some alterations were observed e.g., increased abundance in Alistipides and Bacteroides in AIEC/Piroxicam group, changes often associated with murine colitis (Gkouskou et al., 2014). When individual caecal short chain fatty acids (SCFA) levels were examined, a general but non-significant increase in propionate and reduction in acetate was observed in the AIEC/Piroxicam group (Figure S1E).

Collectively the data showed that AIEC and/or Piroxicam treatment only had modest effects on gut microbiome composition and SCFAs in contrast to other colitis stimuli (Gkouskou et al., 2014).

### Caspase 8 and NLRP3 inhibitors ameliorate NSAID induced epithelial alterations and inflammation in AIEC colonised mice

To further investigate the role of NLRP3 and Caspase-8 in our model, *IL10* ^*-/*-^ mice were administered the selective NLRP3 inhibitor, MCC950 (Xiao et al., 2022), and the Caspase-8 inhibitor Z-IETD-Fmk (Silva et al., 2005), with the treatment of mice starting before AIEC and piroxicam exposure (Figure 2A). Both inhibitors significantly reduced the disease activity index and colon weight of animals administered AIEC and Piroxicam (Figure 2B-C). These inhibitors improved the loss of epithelial integrity, as shown by recovery of colonic *Zo1* and *Muc2* gene expression (Figure 2D). Similarly, histology analysis revealed a significant recovery of epithelial structure represented by less elongated crypts and reduced presence of crypt abscesses (H&E staining) and an increase in mucus-secreting goblet cells (Alcian blue/PAS staining) (Figure 2E-F). Furthermore, inhibition of NLPR3 or Caspase-8 reduced the infiltration of inflammatory cells into the lamina propria (Figure 2F). By flow cytometry, we also observed that the caspase-8 inhibitor significantly reduced activated T-cells (CD4 ^+^ CD69 ^+^) from the spleen while NLRP3 inhibition reduced both activated T-cells and monocytes/macrophages (CD11b ^+^ CD14 ^+^ and CD11b ^-^ CD14 ^+^) while M2 macrophages (CD206 ^+^CD163 ^+^) were increased (Figure 2G and Figure S2A-C). A similar trend in activated T-cells was seen in the mesenteric lymph nodes treated with either of the inhibitors (data not shown). These changes in immune cell populations were accompanied by a reduction in colonic gene expression of acute inflammatory cytokines (*Il6, Cxcl2, Inos)*; macrophage activation markers (*Tnfa, Ccl2*) and T-cell markers (*Ccl5, Cxcl10, Il17a, and Ifng*) (Figure 3A); all of which were initially altered in the AIEC/piroxicam group (Figure 3A, 1D-E and S1B-C). Similarly, a reduction in protein levels of IL-6 and IFN-γ was also observed in mice treated with either inhibitor compared to AIEC/piroxicam mice (Figure 3B). Together, these findings suggest that activation of the NLRP3 inflammasome and Caspase-8 induce epithelial cell alterations leads to inflammation due to the presence of activated T-cells and macrophages in NSAID-exposed *IL10* ^*-/-*^ mice pre-colonised with AIEC.

**Figure 2.**
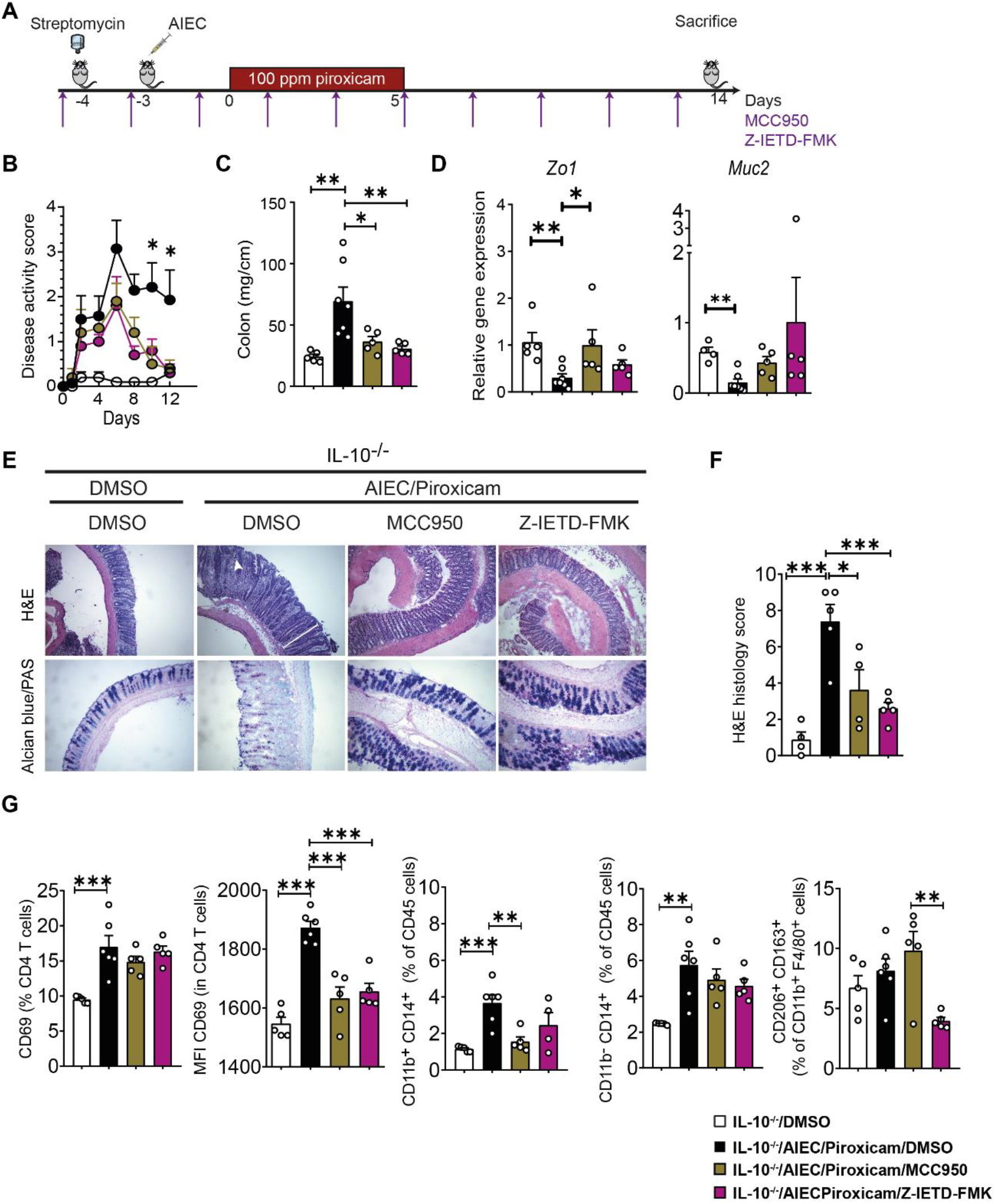
Caspase-8 and NLRP3 inhibitors improve AIEC induced epithelial and immune alterations in IL10 ^-/-^ mice fed with piroxicam. (A) Study design as per Figure 1A. Mice were injected intra-peritoneally with NLRP3 inhibitor (MCC950, 20mg/kg) and Caspase-8 inhibitor (Z-IETD-FMK, 10mg/kg) starting at day -4 and every second day as indicated, followed by euthanasia at day 14. (B) Colon weight. (C) Disease activity index. (D) RT-qPCR of colonic epithelial genes. (E) Representative Haematoxylin and Eosin and alcian blue (AB)/PAS staining of distal colon sections. In AB/PAS staining, Goblet cells are stained in dark purple colour. White arrow indicates crypt abscess and white line indicate crypt hyperplasia (F) Histology score. n=4-7/group. (G) Isolated spleen T-cells (CD4 and CD69) and macrophages (CD11b, CD14, CD163, CD206) were immunophenotyped by Fluorescence-activated cell sorting (FACS) after gating on CD45 (Figure S2A-C) and expressed as MFI or percent of specific cell populations. n=5-6/group. Significance was determined using one-way ANOVA with Bonferroni or Kruskal-Wallis test with Dunn’s multiple comparison test, respectively. * p < 0.05; ** p <0.01; ***p <0.001.

**Figure 3.**
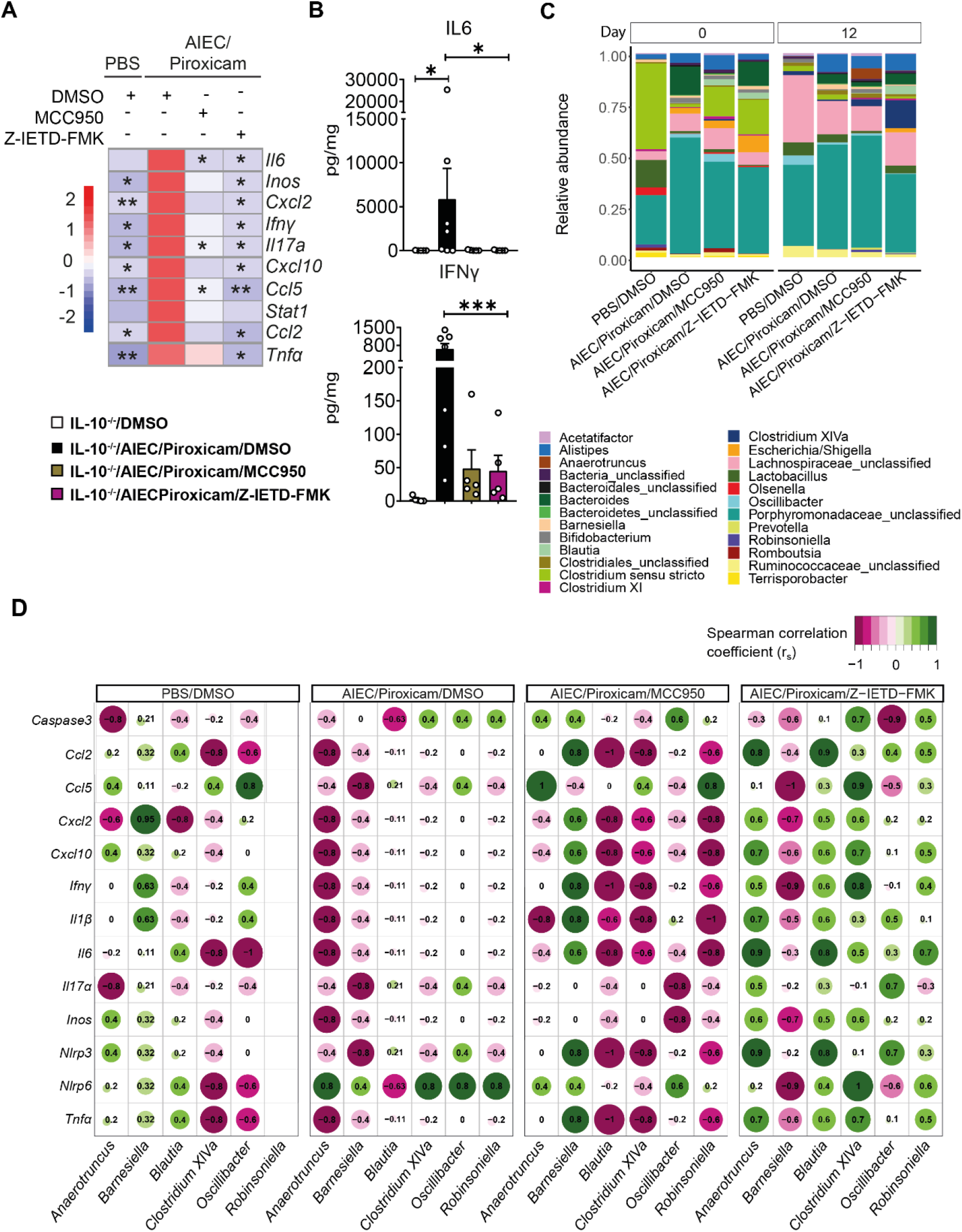
Caspase-8 and NLRP3 inhibitors have a differential effect on host-microbe interaction in AIEC-colonised IL10 ^-/-^ mice treated with piroxicam. (A) Heat map of colonic gene inflammatory expression. AIEC-piroxicam/DMSO treated group was used as control for statistical analysis. (B) Protein levels of colonic cytokine and chemokines n=5-7/group. Data are presented as mean ± SEM. (C) Faecal samples were collected for 16S rRNA analysis at day 0 (upon piroxicam feeding) and 14 (end of trial). Data is presented as faecal bacteria relative-abundance at genus-level. (D) Spearman correlation between host inflammatory gene expression and specific microbial genera at day 14. n=4-9/group. Significance was determined using one-way ANOVA with Bonferroni or Kruskal-Wallis test with Dunn’s multiple comparison test, respectively. * p < 0.05; ** p <0.01; ***p <0.001.

### Differential microbiome-host interactions are associated with effects of NLRP3 and Caspase-8 inhibition

Next, we investigated the effects of the NLRP3 (MCC950) and Caspase-8 (Z-IETD-FMK) inhibitors on bacterial composition in faecal samples collected on day 0 and 12 from *IL10* ^*-/-*^ mice with colitis induced by AIEC/Piroxicam. No significant changes were detected on 16S rRNA alpha-diversity between the different groups or over time (Figure S2D),

When we investigated the variance associated with sample beta-diversity differences (Figure S2E), the variance attributable to days post piroxicam was 12.3%, while the treatment grouping variable was associated with a 11.4% variance. The interaction of these two variables was 8.8% (p-value = 0.011).

To avoid over-cluttering of the compositional bar plot of AIEC/Piroxicam inflamed mice versus Caspase-8/NLRP3 inhibitor groups, only taxa with a total aggregated relative abundance of 0.2% are shown (Figure 3C). Visually, the 16S compositions at day 0, showed a higher abundance in *Clostridium sensu stricto* in the inhibitor treated groups, while at day 12, *Clostridium* cluster XIVa abundance are seen in mice treated with both inhibitors. However, these did not reach statistically significant after fdr correction, potentially due to the small group size comparisons (Figure 3C).

Interestingly, the rank order Spearman correlations of microbial taxa versus inflammatory markers showed contrasting results with regards to the inhibitor treatment (Figure 3D). For putative *Anaerotruncus, Barnesiella, Blautia, Clostridium XIVa, Oscillibacter*, and *Robinsoniella*, the correlation coefficients were almost exactly opposite for NLRP3 (MCC950) and Caspase-8 (Z-IETD-FMK) inhibitors. These microbial correlation differences occur despite both inhibitors similarly dampening the colonic epithelial and inflammatory gene expression profile (Figure 3A).

When the individual caecal SCFA levels were examined, a non-significant increase in propionate was observed in the Caspase-8, and NLRP3 treated groups compared to AIEC/Piroxicam group (Figure S2F).

Overall, this data indicates that NLRP3 and Caspase-8 inhibitors target specific microbiome-host interactions without provoking dramatic alterations on microbial taxa.

### Caspase-8 and NLRP3 inhibitors block inflammasome activation, apoptosis and pyroptosis in AIEC colonised and NSAID exposed mice

AIEC/Piroxicam combination in *IL10* ^*-/-*^ mice resulted in activation of the inflammasome, apoptosis and pyroptosis (Figure 1G). When assaying the expression of genes associated with inflammasome activation and apoptosis in mice administered either the NLRP3 or Caspase-8 inhibitors, no changes were observed at the gene expression levels of *Caspase 1* and *Caspase 8*, while *Il1b, Nlpr3* and *Caspase 3* were significantly reduced when compared to AIEC/Piroxicam group (Figure 4A-B). In support of these findings, western blot analysis showed decreased expression of the inflammatory active cleaved Caspase-1, IL-1β and IL-18, the pro-apoptotic makers - cleaved Caspase-3 and PARP, and the pyroptotic markers – cleaved Caspase-1, Caspase-11 and GSDMD, in the inhibitor-treated groups compared to AIEC/Piroxicam combination group (Figure 4C). As expected, both the NLRP3 inhibitor (MCC950) and the Caspase-8 inhibitor (z-IETD-fmk) reduced the expression of their own target proteins, and both inhibitors decreased the expression of the same biochemical markers of inflammasome activation, apoptosis and pyroptosis (Figure 4D). Interestingly, western blot analysis indicated that both Caspase-8 and NLRP3 can potentially cross-regulate each other. Caspase-8 inhibition reduced NLRP3 both at the gene and protein levels, while NLRP3 inhibition reduced the expression of active cleaved Caspase-8. Several studies have reported that Caspase-8 may act upstream of NLRP3, by inducing NLRP3 activation via GSDMD-mediated K ^+^-efflux (Chi et al., 2014). In order to confirm the involvement of Caspase-8 and its interaction with NLRP3, we assessed the expression of N4BP1, a downstream substrate of Casaspe-8 (Gitlin et al., 2020). As previously shown in Figure S1D, N4BP1 was cleaved in AIEC/Piroxicam combination group, but not in the NLRP3 inhibitor group (Figure 4D), suggesting a role of NLRP3 in Caspase-8 regulation in our model. In line with our expectations, no N4BP1 cleavage was detected in Caspase-8 inhibitor group (Figure 4D), confirming Caspase-8 inhibition. The downregulation in *Nlrp3* gene expression observed in Casapse-8 inhibitor group could be explained by the regulation of the NF-kB by Caspase-8.

**Figure 4.**
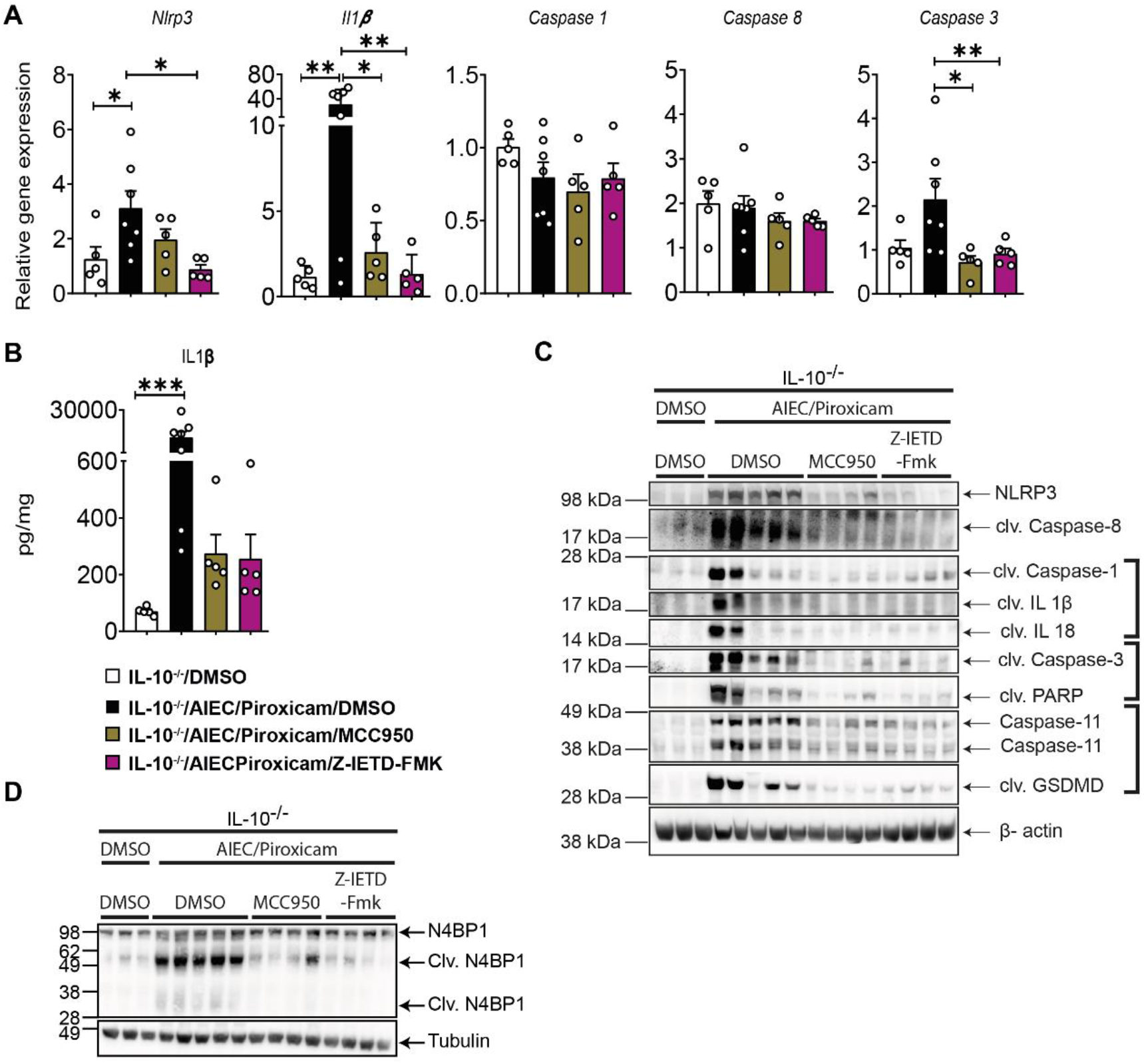
Caspase-8 and NLRP3 inhibitors reduced NLRP3 inflammasome, apoptosis and pyroptosis in AIEC colonised IL10 ^-/-^ mice after piroxicam feeding. (A) RT-qPCR of colonic NLRP3 inflammasome, and caspases. (B) IL1β protein expression. (C) Western blot of Caspase-8, inflammasome and cell death markers. (D) Western blot of N4BP1. n=3-7/group. Data are presented as mean ± SEM. Significance was determined using one-way ANOVA with Bonferroni or Kruskal-Wallis test with Dunn’s multiple comparison test, respectively. * p < 0.05; ** p <0.01; *** p <0.001.

## DISCUSSION

In this study, we examined the hypothesis of whether the presence of an IBD-associated pathobiont (AIEC) (Palmela et al., 2018, Shawki and McCole, 2017) would support the NSAID-worsening symptoms often attributed in patients with IBD. AIEC is a gut pathobiont associated with the pathogenesis of IBD. Consistent with our hypothesis, we show that AIEC colonisation determined the severity of NSAID-triggered inflammation in *IL10* ^*-/-*^ mice which was accompanied by activation of the NLRP3 inflammasome, Caspase-8, apoptosis and pyroptosis, promoting inflammation and cell death.

For our studies, we modified a previously reported model of piroxicam-accelerated colitis (PAC) (Holgersen et al., 2014) by pre-colonising *IL10* ^*-/-*^ mice with AIEC followed by 5 days of piroxicam exposure. Using this design, we showed that piroxicam only induced colitis in mice pre-colonised with AIEC, but not in mice colonised with AIEC alone or treated with NSAID alone. Like the PAC model, the inflammation in the AIEC/piroxicam combination group was accompanied by elevated levels of pro-inflammatory cytokines (e.g., IL-6, TNFα, IFNγ) and infiltration of activated immune cells including T-cells and macrophages. Changes in epithelial structure (altered tight junction proteins, elongated crypts, crypt abscesses and goblet cell loss) in the colon of the *IL10* ^*-/-*^ mice was also observed in these mice. In contrast to the PAC model (Holgersen et al., 2014), the conditions were optimised [i.e., a lower dose of piroxicam (100 vs 200 ppm) and fewer days of feeding (5 vs 9 days) were used] to reduce their adverse effects on animal health after antibiotic treatment and AIEC colonisation (data not shown). In the PAC model, ulcerations were detected in severe cases, while we seldom noted ulcerations in the AIEC/Piroxicam combination group. Alterations in the faecal microbiota in AIEC/Piroxicam group were subtle compared to untreated mice and other pre-clinical models such as the dextran sodium sulphate (DSS)-model (Gkouskou et al., 2014). Changes in the experimental design and the less severe impact on epithelial damage in our model compared to, e.g., DSS-model (Melgar et al., 2005b) might have contributed to the subtle microbial alterations.

Previous studies identified that NLRP3 inflammasome-derived IL1β release plays a critical role in NSAID-induced enteropathy (Higashimori et al., 2016) and treatment with NLRP3 inhibitors such as MCC950 ameliorates colitis (Itani et al., 2016, Perera et al., 2018). IL-10 produced from macrophages was also reported to be a negative regulator of NLRP3 when induced by different stimuli (Zhang et al., 2014).

Caspase-8 was activated in AIEC/Piroxicam combination group. Previous studies show that epithelial deletion of Caspase-8 promoted DSS-induced colitis, associated with increased levels of RIP3 and reduced number of anti-microbial Paneth cells, alluding to Caspase-8 regulation of necroptosis in the pathogenesis of colitis (Gunther et al., 2011). However, no changes in necroptotic cell death markers such as RIPK1, RIP3 or MLKL were observed in our study in Caspase-8 inhibitor group.

The small molecule inhibitors targeting NLRP3, and Caspase-8 rescued the inflammatory phenotype observed in the AIEC/piroxicam group, indicating an indispensable role of Caspase-8 and NLRP3 in AIEC/NSAID-induced inflammation and in IBD-pathogenesis. Interestingly, NLRP3 inhibition targeted the inflammatory response in macrophages by increasing the number of CD206 ^+^ CD163 ^+^ M2 macrophage in the spleen and reducing M1 gene expression profile in the colon, which is in line with a previous study in a model of valve stenosis and calcification (Lu et al., 2022). No increase in M2 cells and a non-significant reduction in macrophages (CD11b+CD14+) were seen in the Caspase-8 inhibitor group, indicating potential differences in the regulation of macrophages by these proteins. Both inhibitors significantly reduced the presence of activated T-cells and associated cytokines such as IFNγ and IL-17α and reduced apoptotic and pyroptotic cell death, which probably contributed to the overall ameliorated phenotype presented by either of the inhibitor-treated groups.

No significant alterations in microbiota composition or individual SCFAs were detected in the inhibitor-treated groups, although an increased abundance of *Clostridium* cluster *XIVa* was seen in both NLRP3 and Caspase-8 inhibitor groups. This is a SCFA producing bacteria, it can regulate Treg cells, and its abundance is generally decreased in IBD (Kim et al., 2014, Takeshita et al., 2016). A similar effect on microbiota composition was reported in a model of experimental autoimmune encephalomyelitis (EAE) after NLRP3-inhibitor (MCC950) administration (Xu et al., 2020). However, no reports were found in the literature on the impact of Caspase-8 inhibitor (Z-IETD-Fmk) or piroxicam treatment in *IL10* ^*-/-*^ mice on the microbiota composition. Although no increase in butyrate was noted in either of the inhibitor treatment groups, a higher induction of propionate and total SCFA was observed, potentially indicating the contribution of this genus to SCFA generation and reduced inflammation. The non-significant findings could also reflect the small group size comparisons, for this reason future studies are warranted to further dissect the changes in microbial taxa following the use of the NLRP3 and caspase-8 inhibitors.

In conclusion, our findings provide evidence and mechanistic insights into how NSAID and an opportunistic IBD-associated gut pathobiont can synergise to worsen IBD symptoms and inflammation. To the best of our knowledge, this is the first time that Caspase-8 has been reported as a potential regulator of AIEC and NSAID-induced colitis. Our data suggests that genetic factors alone are not enough to trigger pathogen-like activity in a pathobiont, and other factors such as environmental (exposure to NSAID) are needed to trigger its pathogenic potential. Further, we showed that targeting either Caspase-8 or NLRP3 could be a potential therapeutic strategy for IBD patients with NSAID-worsened inflammation. We propose that future clinical studies focusing on the role of NSAID in IBD should consider the AIEC colonisation status as one of the factors influencing the inflammatory phenotype.

One of the limitations of this study is the lack of information on which cell subset contributes to the activation of the NLRP3 inflammasome, Caspase-8 and biochemical markers of apoptosis and pyroptosis. To mechanistically validate the findings from our model, we used human and murine epithelial cell lines (HT29, DLD1, CMT96) and primary cells including murine intestinal crypts and bone-marrow derived macrophages. However, none of these cell systems responded to *in vitro* activation with piroxicam, although previous studies have shown piroxicam-induced cell death in HCT116 and CaCo-2 cells (Somchit et al., 2009). Keeping these limitations in mind, future studies are needed to further investigate the cross-regulation of Caspase-8/NLRP3 axis. Especially the generation of conditional knock-out mice for Caspase-8 and NLRP3 on an IL-10 background would be highly relevant to pinpoint the cell specificity of Caspase-8/NLRP3 axis activation. Multiple staining for co-localisation of ASC, Caspase-8 and NLRP3 on colon sections would also provide further information on cell-specific activation.

## ACKNOWLEDGMENTS

This work and the authors were supported by a Science Foundation Ireland (SFI) Research Centre awards SFI/12/RC/2273-P1 and SFI/12/RC/2273-P2 to APC Microbiome Ireland and SFI Professor award grant number SFI/15/RP/2828 to SA-E. GS-G is a recipient of a Government of Ireland Postgraduate Scholarship (grant GOIPG/2019/4528). We acknowledge and thank the APC Microbiome Ireland Flow Cytometry Platform and the support of the staff at the BSU Annex, UCC.

## AUTHOR CONTRIBUTIONS

RS – conceived, designed and carried most of the experiments, analysed related data, wrote the initial manuscript and edited the paper; VR – designed, carried out and analysed flow cytometry experiments and executed animal experiments; SRS, AC – analysed and graphed microbiota analysis; GS-G, NH, T-DS – carried out the animal experiments, dissected tissue and cells and staining of cells for flow cytometry; LD, CH – supervised microbiota analysis and edited paper; KN, FS, SA-E – reviewed and edited paper; SM – conceived, designed and supervised the research and edited the paper. FS, SA-E, SM – funding acquisition. All authors contributed to the article and approved the submitted version.

## FIGURE LEGENDS

**Figure S1.**
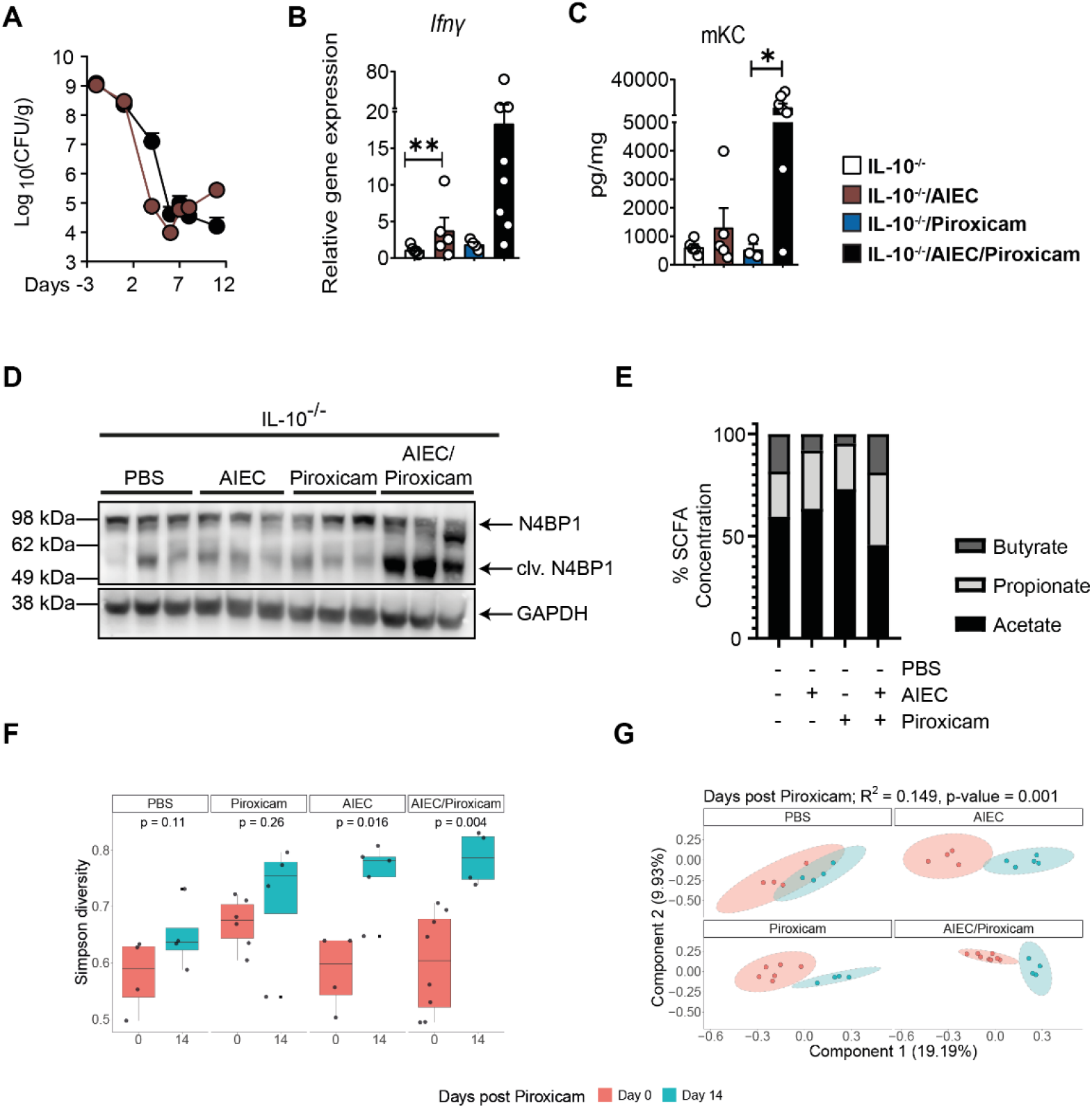
AIEC colonisation and piroxicam treatment subtly altered the gut microbiota composition. (A) Faecal AIEC colonisation. n=5-8/group. (B) RT-qPCR of colonic *Ifng*. (C) Colon mKC protein. (D) Western blot for N4BP1. (E) Caecal SCFA levels. (F-G) Faecal samples were collected for 16S rRNA analysis at days 0 and 14. (F) Simpson diversity graphed by treatment. (G) PCoA analysis. n=4-9/group. For (A-D) data are presented as mean ± SEM. Significance was determined using one-way ANOVA with Bonferroni or Kruskal-Wallis test with Dunn’s multiple comparison test, respectively. * p < 0.05; ** p <0.01, *** p <0.001.

**Figure S2.**
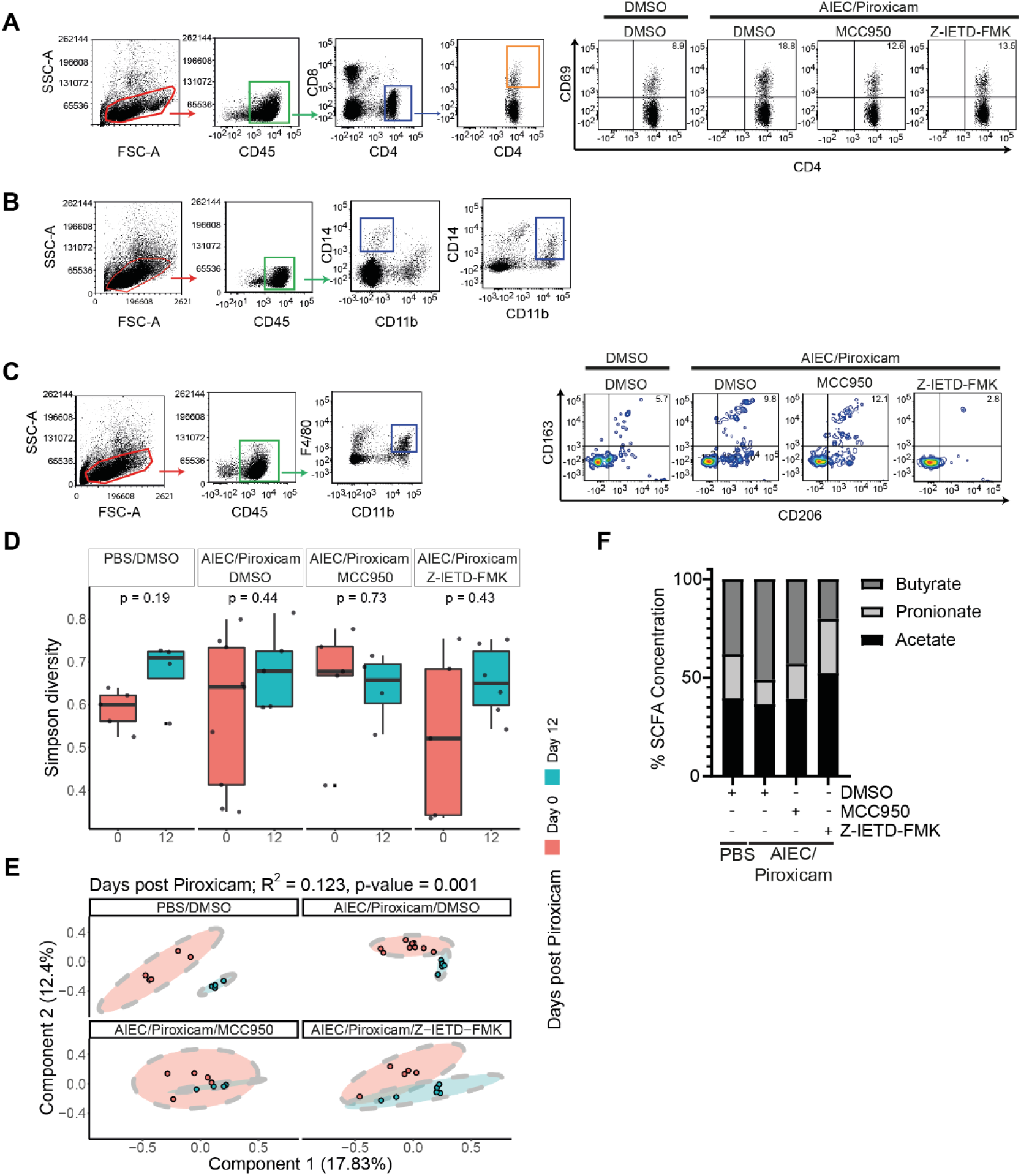
Impact of Caspase8 and NLRP3 inhibition on T cells and macrophage populations and microbiota composition. (A-C) Gating strategy. Spleen activated T-cells (CD4+ CD69+) and macrophages (CD45+ CD11b+ CD14+; CD45+ CD11b-CD14+; CD45+ CD163+ CD206+ M2 macrophages) were immunophenotyped by flow cytometry. (D) Simpson diversity graphed by treatment. (E) PcoA analysis. (F) Caecal SCFA levels. n=4-7/group. Data are presented as mean ± SEM. Significance was determined using one-way ANOVA with Bonferroni or Kruskal-Wallis test with Dunn’s multiple comparison test, respectively. * p < 0.05; ** p <0.01; ***p <0.001.

## METHODS

### Bacterial strain and growth

Adherent-invasive *E. coli* HM605 stain was previously isolated from colonic biopsy of CD patients (Martin et al., 2004) and was provided by Dr David Clarke, UCC. For mouse oral gavage, *E. coli* HM605 was grown by culturing a single colony in 5 millilitre (mL) Luria Bertani (LB) Broth (Cat#L3022, Sigma-Aldrich) with 50 µg/mL ampicillin (Cat#A9518, Sigma-Aldrich) at 37 °C overnight without shaking. The next day, the overnight culture was washed twice with PBS (Sigma) by centrifugation at 500 g for 10 min. After the second wash, the bacterial pellet was re-dispersed in PBS supplemented with 10 % 1M NaHCO _3_ (Cat#S5761, Sigma-Aldrich) with the final OD _600_ of 5. Each mouse was gavaged with 200 μml of bacterial culture with an OD _600_ of 5.

### Mice

IL-10 ^-/-^ mice (B6.129P2-Il10tm1Cgn/J) were acquired from Charles River Laboratories, UK and bred at the Biosciences unit Annex at University College Cork. All animals were housed in individually ventilated cages (IVCs, OptiMICE, UK), with a controlled environment (20–22 °C, 12 hours (hr) light:dark cycle) and given food and water *ad libitum*. All animal studies were designed with consideration for the three Rs (Replacement, Reduction and Refinement) and were approved by the Animal Experimentation Ethics Committee (AEEC; application #2020/007) of University College Cork and by Health Products Regulatory Authority (HPRA, project nr AE19130/P138).

To establish a model of AIEC-colonisation and piroxicam induced colitis, male IL-10 ^-/-^ mice (8-16 weeks) were randomised into four groups, PBS (vehicle), AIEC only, piroxicam only and AIEC and piroxicam combination. To establish colonisation of AIEC, IL-10 ^-/-^ mice were given streptomycin (5 g/L, Cat#S9137, Sigma-Aldrich) in the drinking water *ad libitum* for 24 hr. After streptomycin treatment, mice were orally gavaged with 10 ^8^ to 10 ^9^ colony forming units (CFU) of Adherent-invasive *E. coli* HM605 in 0.2 mL/mouse. Three days after AIEC infection, mice were given 100 parts per million (ppm) of piroxicam (Sigma) homogenised in regular chow (Envigo, UK) for up to 5 days followed by 9 days of regular chow. At day 14, when mice were sacrificed by cervical dislocation and intestinal tissue was collected for further analysis (Figure 1Ay). Faecal samples were collected at day 0, 6 and 14 for 16S rDNA analysis.

For the inhibitor experiment, the NLPR3 inhibitor MCC950 (20mg/kg, Cat#HY-12815A, MedChemExperss) and caspase-8 inhibitor Z-IETD-FMK (10mg/kg, Cat#HY-101297, MedChemExperss) were diluted in DMSO and injected, intraperitoneally, in a 0.2mL volume, every other day starting from day -5 (Figure 2A) followed by streptomycin treatment, AIEC gavage and piroxicam feeding as outlined above. Control mice were injected with DMSO in the same manner as the inhibitors. Mice were sacrificed on day 14 (Figure 2A). Intestinal tissue, spleen, mesenteric lymph nodes (MLNs) and caecal samples and blood were collected at sacrifice for further analysis. Faecal samples were collected at days 0 and 12 various time points during the study for 16S rDNA analysis.

### Collection of samples

Colons were excised, opened longitudinally, and washed in PBS. Colon length was recorded, and colon was divided in 2 pieces with 3 cm of distal colon (weighed) and divided in two longitudinally pieces; one was snap frozen in liquid nitrogen for western blot analysis, and the other section was rolled, embedded in optimum cutting temperature (OCT, Cat#4583, Tissue-Tek™) compound and snap frozen in liquid nitrogen for histological analysis. For RNA isolation and RT-qPCR, a 0.5 cm piece of the most distal colon was stored in RNAlater (Cat#3335402001, Roche) for 24 hr at 4 °C and snap frozen in liquid nitrogen. The spleen and mesenteric lymph nodes were removed and processed for isolation of cells (see below). Faeces and caecal content were collected, snap frozen in liquid nitrogen for 16S rDNA analysis. All collected samples were stored at -80 °C until needed.

### Haematoxylin and Eosin and PAS staining and scoring

OCT-embedded distal colon sections were cut in Leica Cryostat (Model) at 3-5mm and fixed in 10% Neutral Buffered formalin, followed by haematoxylin and eosin (H&E) staining as previously reported (Hall et al., 2013b). For detection of mucus producing cells, sections were fixed in 100% Iso-propyl alcohol, stained with Alcian blue and periodic acid-Schiff (PAS) previously reported (Las Heras et al., 2019). H&E-stained samples were scored according to (Holgersen et al., 2014) with some modifications. Briefly, inflammation (0-4), hyperplasia (0-4) and extent of area involved (0: 0-10%; 1: 10-25%; 2: 25-50%; 3: 50-75%; 4: 75-100%) were scored individually resulting in a total score of 12. No areas of ulcerations were observed in the sections. Pictures were taken with Leica DMLB microscope at 10x zoom.

### Bacterial counts

To enumerate the bacteria in the stool samples, faecal pellets were weighed and homogenised in PBS at a concentration of 0.1 mg/mL. Homogenised samples were plated onto LB agar with 50 µg/mL ampicillin. After culturing at 37 °C overnight, bacterial counts were recorded.

### Real time qPCR analysis

Total RNA from the colonic tissue was isolated using the Qiagen RNeasy kit (Qiagen) following the manufacturer’s protocol and quantified using a NanoDrop Spectrophotometer (ND1000). The RNA was treated with Turbo DNA-free kit (Invitrogen), to remove genomic DNA, following the manufacturer’s instructions. 1 μg of RNA was reverse transcribed using Transcriptor Reverse Transcriptase from Roche. The qPCR was carried out using a Roche LightCycler 480 instrument and SensiFAST No-ROX mix. The Ct values obtained were compared using 2-ΔCt. The expression of genes was normalised to that of β-actin. Primers and probes were designed using Universal Probe Library Assay Design Center (https://www.roche-applied-science.com/sis/rtpcr/upl/adc.jsp; Roche Applied Science). Primer and probe sequences of all genes analysed are summarised in Table 1.

### Meso Scale Discovery (MSD) multiplex assay for cytokine for protein quantification

Colonic tissue was homogenised as previously described (Hall et al., 2013a). Briefly, colon tissue was homogenised in 350 µl of lysis buffer (40 ml of PBS, 10%FCS, 2 Complete Protease Cocktail Inhibitor Tablets (Roche, Mannheim, Germany). Each sample was subject to 3 rounds of homogenisation using a MagnaLyser (Roche) at 6,000 rpm for 15 seconds, placing on ice for 30 seconds between each round, followed by centrifugation at 10,000 g for 10 minutes at 4 °C. The supernatants/homogenates were aliquoted into fresh 1.5ml tubes and stored at -80 °C until used. 25 µl of lysates per sample were used in MSDmeso scale discovery U-plex assay to quantify the cytokines IFNγ, IL1β, IL6 and mKC according to the manufacturer’s instructions. Briefly, U-plex 10 assay plate was coated with linker-coupled biotinylated capture antibody overnight at 4 °C. Next day, the plate was washed thrice with PBS containing 0.1% tween-20 (PBST). After washing, samples and standards were added to the plate and incubated at room temperature (RT) for 1 hour with shaking, followed by three washes with PBST. 50 µL of secondary detection antibodies solution were added for 1 hour at RT with shaking following PBST washes. After secondary incubation, the plate was washed thrice with PBST, followed by 150 µL of read buffer. The plate was read immediately after adding read buffer using MESO QuickPlex SQ 120. Data is graphed as cytokine levels are expressed as pg cytokine/mg colonic tissue.

### Western blot analysis

Western blot was performed as described with some modifications (Saiz-Gonzalo et al., 2021). Briefly, colonic tissue samples were homogenised in tissue lysis buffer (50 mM NaCl, 50 mM NaF, 50 mM Na _4_P _2_O _7_, 5 mM EGTA, 5 mM EDTA, 2 mM Na3VO4, 10 mM HEPES, 1% Triton X-100, pH 7.4) supplemented with 1 × Halt Protease and Phosphatase Inhibitor Cocktail (cat no. 78440, Thermo Scientific), 0.02 mg/ml RNase, 0.2 mg/ml DNase, 0.01 mM Na _3_VO _4_, 0.005 mM Na _4_P _2_O _7_, 0.01 mM β-glycerophosphate. Lysate was cleared by centrifugation for 15 min at 14,000 rpm and 4 °C, and protein concentration was measured using Pierce BCA Protein Assay Kit (cat no. 23225, Thermo Scientific). Forty to eighty micrograms of protein samples were denatured in 1 × Bolt LDS Sample Buffer (Invitrogen) supplemented with 1 × Bolt Sample Reducing Agent (Invitrogen) by heating for 10 min at 75 °C. Samples were separated on Bolt 4–12% Bis-Tris Plus Gels (Invitrogen) and transferred to a PVDF membrane (cat no. IPVH00010, Millipore) using the Mini Blot Module (Invitrogen). The membrane was blocked with 5% TBS with 0.1% Tween-20 (TBST) for 1 hour, at RT and incubated with the primary antibody overnight at 4 °C. Next day, the membrane was washed with TBST and probed with the secondary antibody for 1 hour, at RT followed by TBST wash and detection with WesternBright Quantum HRP substrate (cat no.

K-12042-D10, Advansta) and LAS-3000 Imager (Fujifilm) and processed with ImageJ software (without gel splicing and brightness/contrast adjustment).

### Flow cytometric analysis and cell sorting

Spleens were removed and single-cell suspensions were prepared as reported with some modifications (Hall et al., 2011). Briefly, to isolate cells from the spleen, the tissue was pressed through a 100 μm cell strainer (Sarstedt, Germany) positioned on a 50 mL tube using the plunger of a 1 mL syringe and washing the strainer with 1X sterile PBS supplemented with 1% FCS. Samples were centrifuged for 5 min at 300 x g and red blood cells were lysed by 10 min incubation on 37 °C with 5 mL of 1 X RBC lysis buffer (eBioscience). Immune cells were then re-suspended in 1 X PBS supplemented with 1 % FCS. Isolated cells were washed three times in PBS supplemented with 1 % bovine serum albumin (BSA, Sigma) and 0.1% sodium azide (Sigma). Non-specific binding of antibodies (Abs) to Fc receptors was blocked by pre-incubation of cells with monoclonal Abs (mAb) 2.4G2 directed against the FcgRIII/II CD16/CD32 (0.5 ng mAb per 10 ^6^ cells). 1 × 10 ^6^ cells were incubated with 0.5 ng of the relevant mAb for 20 min at 4 °C and washed twice. mAbs used in this study are listed in the key resource table. Data were analysed using FCS Express V5 software (De Novo). Cells were analysed using the three laser (405nm, 488nm, 460nm) BD Celesta FACS Analyzer. The forward narrow angle light scatter was used as an additional parameter to facilitate the exclusion of dead cells and aggregated cell clumps. A forward scatter height (FSC-H) vs forward scatter area (FSC-A) density plot was used to exclude doublets. Then, FSC-A vs side scatter area (SSC-A) density plot was used to identify cells of interest. For T cells analysis, CD4 vs CD8 dot plots were obtained after gating cells with CD45 expression. For macrophage analysis, CD11b vs CD14 and CD11b vs F4/80 dot plots were obtained by gating on cells with CD45 expression. CD206 vs CD163 dot plots were obtained after gating on CD11b ^+^F4/80 ^+^ cells.

### DNA extraction, 16S rRNA gene sequencing, and gut microbiota analyses

DNA from faecal samples were extracted using a QIAamp Fast DNA Stool Mini Kit (Qiagen cat no. 51604) following the manufacturer’s instructions. Amplicon sequencing of the 16S rDNA V3-V4 region of bacterial communities was performed as previously described (Ryan et al., 2020), utilising 2×250bp paired-end Illumina MiSeq chemistry (Genewiz; Leipzig, Germany). Raw sequencing reads were processed using an in-house 16S rRNA processing pipeline, employing USEARCH (64 bit; v 8.1). Briefly, paired-end reads were merged and filtered using a <0.5 expected error rate per nucleotide and total length. Reads were dereplicated, and singletons removed, following the trimming of the forward and reverse primers (“-stripleft 17” and “-stripright 21”, respectively). Operational taxonomic units (OTUs) were clustered at 97% identity, and reference-based chimera removal was performed using UCHIME. OTUs were assigned taxonomic information by aligning reads to the RDP Gold database using the RDP Classifier (v 2.12).

The 16S rRNA reads assigned taxonomic information was converted into a count matrix and imported into R Studio (v 3.6.1). The 16S data was analysed using the metadata accompanying the relevant mouse trials and the RT-PCR experiments (Supplementary Data). The 16S rRNA OTU reads aligned per OTU were converted into relative abundances using the “funrar” package (Grenié et al., 2017). Dataframes and matrices were manipulated as necessary using the “reshape2” package (Wickham, 2007). Publication quality images were generated using the “ggplot2” and “ggpubr” packages (Wickham, 2016, Kassambara, 2019). Boxplots represent the standard Tukey representation, with boxes representing the 25 ^th^, 50 ^th^ (median) and 75 ^th^ interquartile range (IQR) percentiles, and the whiskers encompassing values within 1.5x the IQR. The specific values of each boxplot are overplotted as opaque grey circular points using the ggplot “geom_jitter” function, whereas boxplot outliers are represented as solid square points.

The alpha- and beta-diversity of samples were calculated using the R packages “vegan” and “phyloseq” (Oksanen et al., 2013, McMurdie and Holmes, 2013). All alpha-diversity values presented use Simpson’s index, while beta-diversity separation was performed using Canberra distances with PCoA ordination. A colour palette for the composition bar plots was obtained through the “pals” R package (Wright, 2019). The log _2_ fold change of epithelial and inflammatory biomarker genes, and 16S rRNA taxa, was performed using the “DESeq2” package after scaling values to integers (Love et al., 2014). Correlations between microbial taxa and epithelial and inflammatory biomarker genes were performed using base R’s “stats” package (Team, 2019), with plots generated using the “corrplot” package (Wei et al., 2017). Individual images were manipulated into their final multipanel display using Inkscape (v 1.1.2).

### Short chain fatty acids (SCFAs)

Analysis of SCFAs was performed as recently reported (Aguilera et al., 2021). Briefly, Caecal samples were weighed and diluted 1:10 (w/v) in sterile HPLC grade water. The SCFA-containing supernatant was filtered through 0.2 µm pore size cellulose acetate membrane (GyroDisc CA; Orange Scientific, Braine-l’Alleud, Belgium) and stored at −20 °C until HPLC analysis. Quantification of SCFAs in caecal samples was carried out using an external calibration standard curves method (Aguilera et al., 2021). Caecal SCFA concentrations were expressed as mean µmol per gram wet weight cecum using the following equation: Caecal SCFA (µmol/g) = [organic acid in caecal contents (mmol/mL) X Vd (ml) × 1000]/wet weight cecum (g), where Vd = total volume of dilution.

### Statistical analysis

GraphPad Prism software was used to perform statistical analyses for all data sets except 16S amplicon data. All data were expressed as mean ± SEM One-way ANOVA with Bonferroni test or Kruskal-Wallis with Dunn’s multiple comparison test was used to test for significant differences between various groups. A p-value <> 0.05 was considered significant.

For analysing the 16S amplicon data of the relevant mouse trials, the following statistical tests were performed. Wilcoxon tests were performed for nonparametric two-group comparisons, with the p-values presented above the relevant alpha-diversity box plots. Statistical significance in the beta-diversity of mice with respect to days post piroxicam, treatment group, or their interaction was assessed using the “adonis” function of the vegan R package, which performs a permutational multivariate analysis of variance (PERMANOVA) test. The variance (R ^2^) and p-values of PERMANOVA tests accompany the relevant PCoA plots. Statistically significant changes in microbial taxa with reference to mice treatment groups were calculated using Wilcoxon tests with Benjamini-Hochberg false discovery rate (fdr) correction. All correlations between microbial taxa and epithelial and inflammatory biomarkers were conducted using Spearman rank correlations.

### Data code and material availability

Raw sequence data from mouse faecal samples have been deposited in the NCBI SRA database under accession number SAMN29132731 to SAMN29132837 for colitis study and SAMN29826455 to SAMN29826520 for the inhibitor study under the Bioproject number PRJNA849757. This paper does not report any original code or unique reagent. Any additional information required to reanalyze the data reported in this paper is available from the lead contact, Silvia Melgar, upon request.

## Reagent and resource

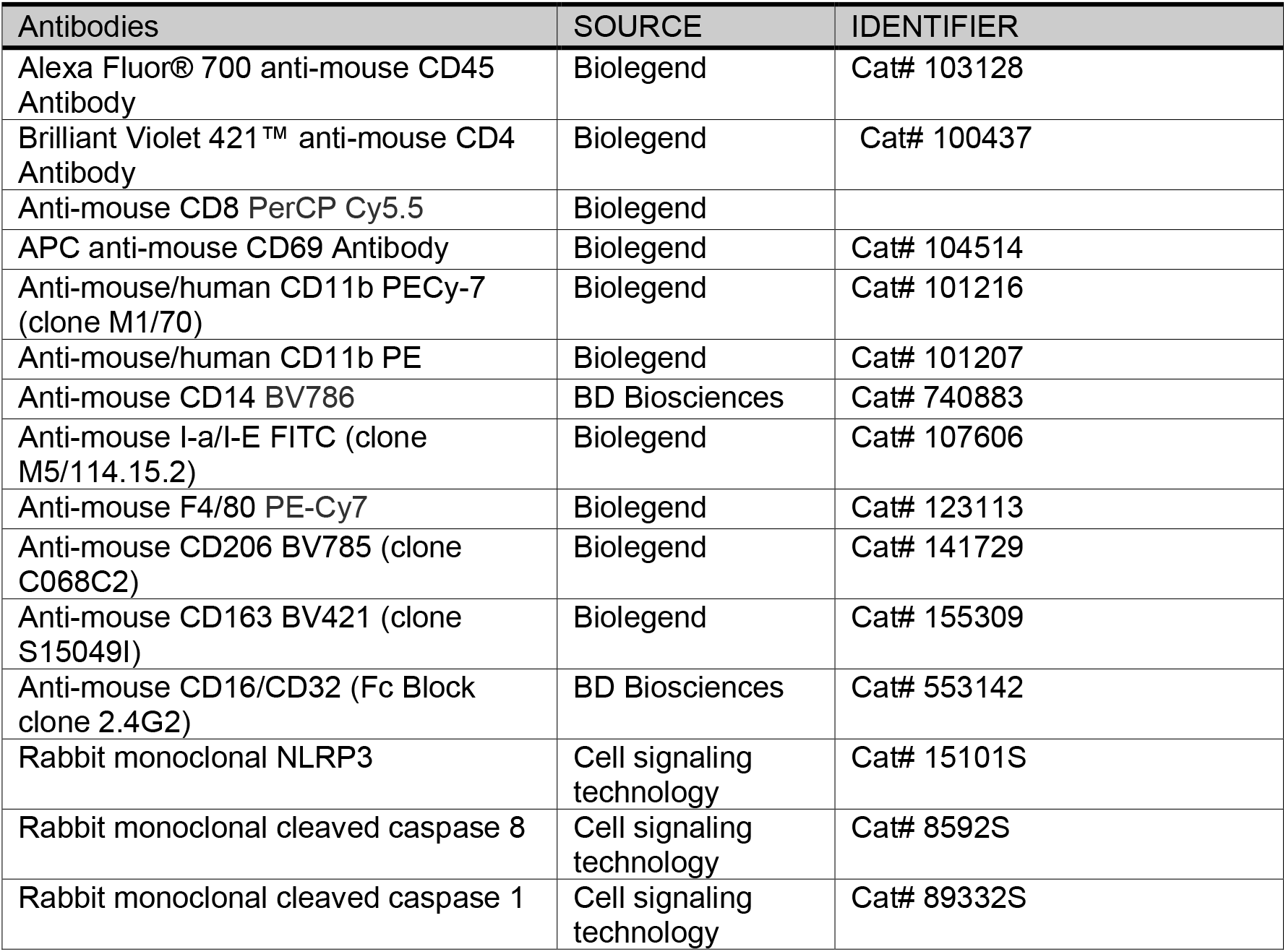
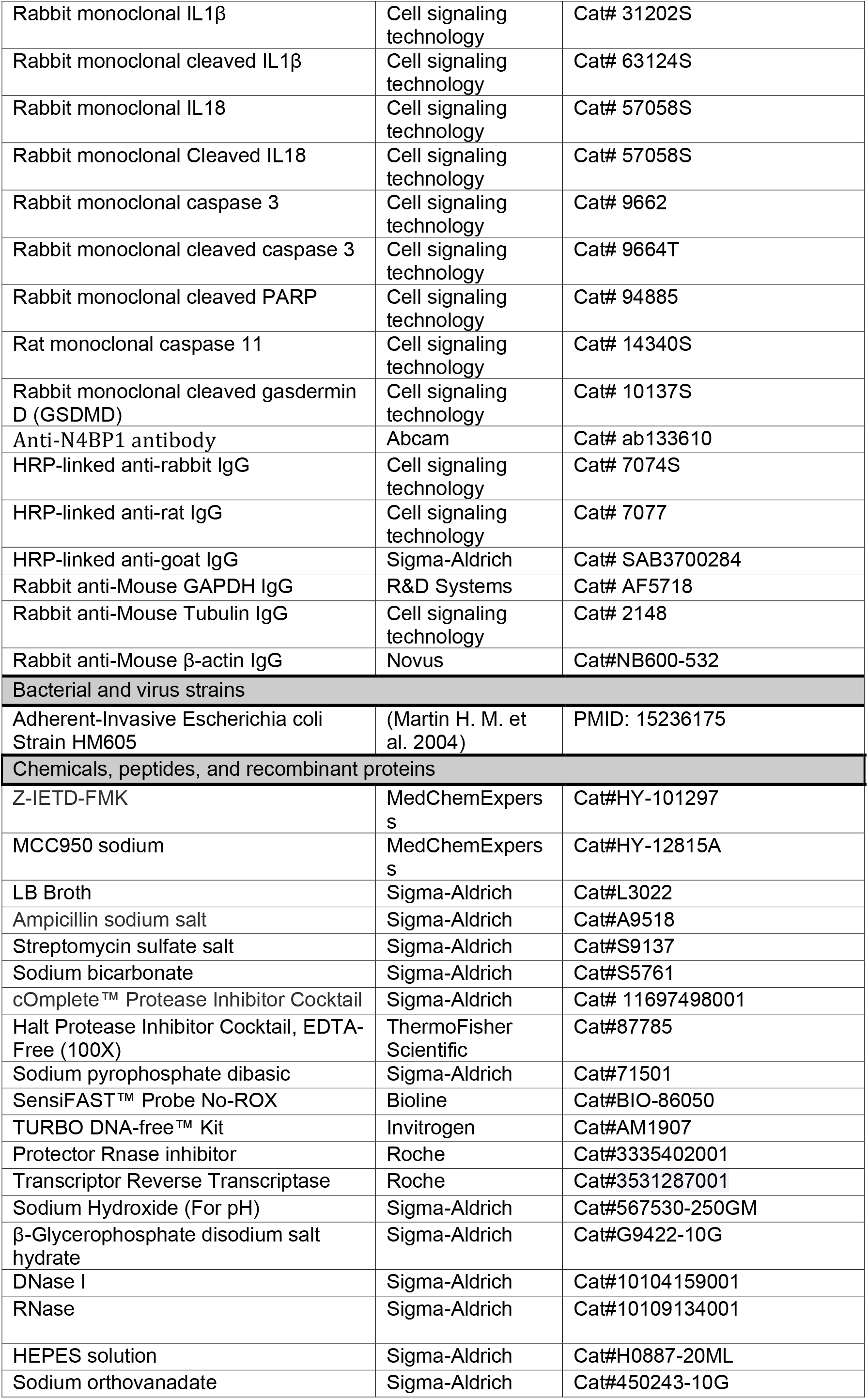
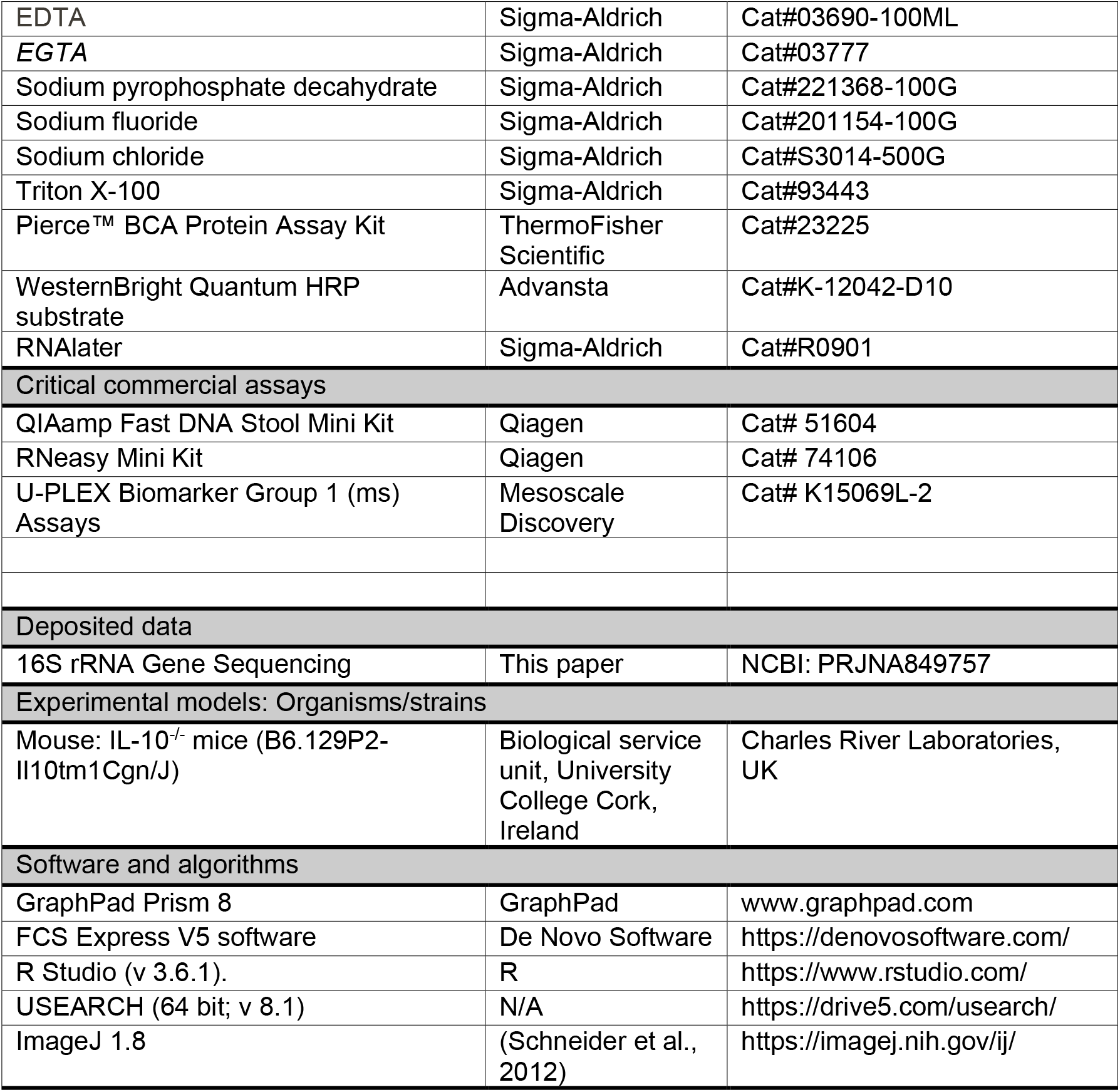

## Primers used in the study

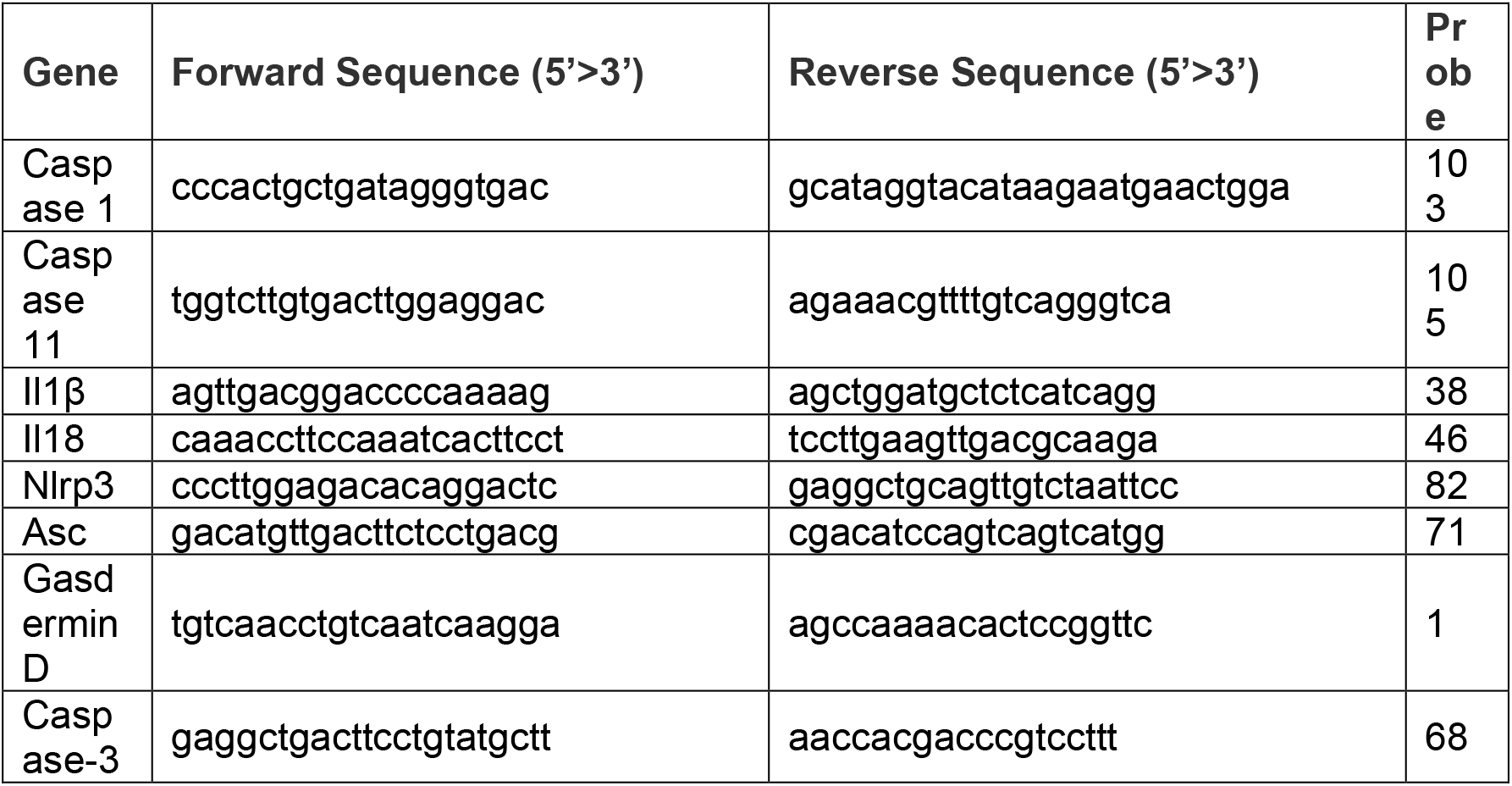
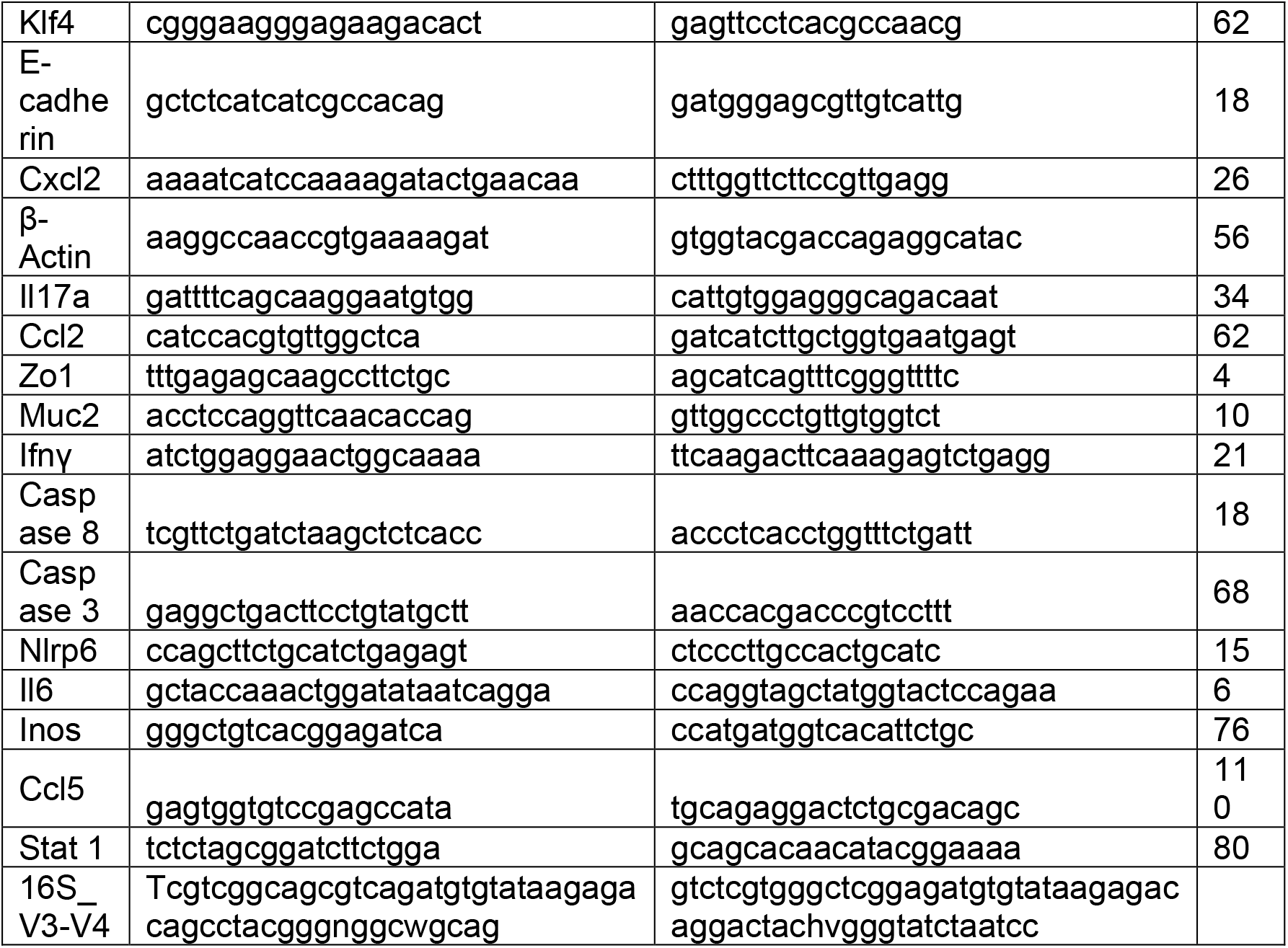

